# Spatiotemporal characterization of periocular mesenchyme heterogeneity during anterior segment development

**DOI:** 10.1101/726257

**Authors:** Kristyn L. Van Der Meulen, Oliver Vöcking, Megan L. Weaver, Jakub K. Famulski

**Affiliations:** Department of Biology, University of Kentucky

**Keywords:** Periocular Mesenchyme, Anterior segment, Anterior Segment Dysgenesis, Neural crest, *Pitx2*, *FoxC1*

## Abstract

Establishment of the ocular anterior segment (AS) is a critical event during development of the vertebrate visual system. Failure in this process leads to Anterior Segment Dysgenesis (ASD), which is characterized by congenital blindness and predisposition to glaucoma. The anterior segment is largely formed via a neural crest-derived population, the Periocular Mesenchyme (POM). In this study, we aimed to characterize POM behaviors and identities during zebrafish AS development. POM distributions and migratory dynamics were analyzed using transgenic zebrafish embryos (Tg[*foxC1b:GFP*], Tg[*foxD3:GFP*], Tg[*pitx2:GFP*], Tg[*lmx1b.1:GFP*], and Tg[*sox10:GFP*] throughout the course of early AS development (24-72hpf). *In vivo* imaging analysis revealed unique AS distribution and migratory behavior among the reporter lines, suggesting AS mesenchyme (ASM) is a heterogenous population. This was confirmed using double *in situ* hybridization. Furthermore, we generated ASM transcriptomic profiles from our reporter lines and using a four-way comparison analysis uncovered unique ASM subpopulation expression patterns. Taken together, our data reveal for the first time that AS-associated POM is not homogeneous but rather comprised of several unique subpopulations identifiable by their distributions, behaviors, and transcriptomic profiles.

## INTRODUCTION

Vertebrate cranial development has benefitted significantly from the evolutionary addition of the multipotent neural crest cells (NCC). In the developing cranial region, migrating NCCs come together with lateral plate mesoderm to surround the developing optic cup and form the periocular mesenchyme (POM) (1–3). POM cells subsequently contribute to the development of the ocular anterior segment (AS) (3–6). The AS, comprising of the cornea, lens, iris, ciliary body, and drainage structures of the iridocorneal angle, is essential for the function of the eye. The AS focuses light onto the retina and maintains intraocular homeostasis.

AS development begins after the establishment of the optic cup, when POM cells migrate into the periocular space between the retina and the newly established corneal epithelium (5, 7). These mesenchymal cells will eventually differentiate into the corneal stroma and endothelium, iris and ciliary body stroma, and the iridocorneal angle, amongst others. Mis-regulation of POM migration and/or function has been associated with congenital blinding disorders under the Anterior Segment Dysgenesis (ASD) label. ASD includes, alone or in combination, corneal opacity, iris hypoplasia, polycoria, corectopia, posterior embryotoxon, juvenile glaucoma, and disorders including Peter’s Anomaly and Axenfeld-Rieger Syndrome (4, 8, 9). These rare autosomal dominant disorders, in addition to ASD phenotypes, also often exhibit systemic issues including dental malformations and craniofacial defects (10–12).

The most common mutations seen in ASD patients involve the transcription factor *pitx2* (Paired-like homeodomain) (11), as well as *foxC1* (Forkhead Box C1) (10, 13–16). Loss of function of either *pitx2* or *foxc1* has been shown to result in ASD phenotypes in mice and zebrafish (11, 13–18). *Pitx2* in particular has been associated with the survival and migration of NCCs, as well as the development of the optic stalk, establishment of angiogenic privilege within the cornea, and craniofacial development (10, 11, 14, 17–21). *FoxC1* and *pitx2* are also known to interact with one another and their expression is regulated by retinoic acid signaling (13, 22, 23). Not surprisingly, mutations in NCC regulatory genes have also been associated with ASD. *FoxD3* (Forkhead Box D3) has been implicated in ASD (24) and is known to regulate early NCC specification, migration and long-term cell survival (25–28). *Sox10* (SRY-Box 10), another key regulator of the NCC population (1, 3, 5, 25, 29), is critical for NCC migration and viability during early development (29). Finally, *lmx1b* (Lim homeodomain) is associated with Nail-Patella syndrome and glaucoma predisposition (30, 31). *Lmx1b* is expressed within the developing cornea, iris, ciliary bodies, and trabecular meshwork of the iridocorneal angle in mice (30, 31) and is important for POM migration in zebrafish (31).

While all these genes have been linked as markers of POM, a molecular mechanism defining AS formation is lacking. Specifically, little is known about when or how POM cells acquire their AS targeting, migrate to appropriate areas, interact with one another or specify into AS structures. Using zebrafish embryos, we sought to investigate and characterize the precise migration patterns and transcriptional profiles of POM cells. In doing so, we catalogued AS distribution, migratory dynamics, and population sizes of several key POM populations throughout AS development. Our findings indicate that AS associated POM is composed of several subpopulations, each identifiable by their distribution, migratory pattern and gene expression profile.

## RESULTS

### POM-associated genes exhibit unique expression patterns during formation of the anterior segment

Work from model organisms as well as clinical investigation into the genetic cause of ASD has identified several genes associated with AS development (32). Critically important are *foxC1* and *pitx2*, but also *lmx1b* and *eya2* as well as NCC-associated regulators *foxD3* and *sox10*. With several genes being implicated in POM migration and identity, we first chose to carefully characterize patterns of their expression during zebrafish ocular morphogenesis (12-72hpf). Whole mount *in situ* hybridization (WISH) using embryos aged 12, 18, 24, 32, 48 and 72hpf revealed that POM-related genes *foxc1a, foxc1b, eya2, foxd3, pitx2, sox10, lmx1b.1* and *2*, display both overlapping and individualized expression patterns within their originating neural crest streams and surrounding the AS (Fig. S1). POM-related genes begin to show expression as early as 12hpf. As the optic cup begins to take shape (18hpf), *foxc1a, foxc1b* and *sox10* expressing POM cells are already visible within the craniofacial space (Fig. S1). At the same time, *pitx2* expression is absent from periocular regions and presents primarily in the lens. By 24hpf we observe periocular expression of all the aforementioned POM associated genes. This pattern persists up to 48hpf for *foxc1a, foxc1b, eya2, pitx2*, and *sox10*. By 72hpf only *foxc1a, sox10*, and *eya2* still exhibit AS associated expression. In spite of the implicated role of *Lmx1b* genes in pathogenesis of the Nail-Patella Syndrome (30, 31), both *lmx1b. 1* or lmx1b.*2* genes do not display classical AS expression patterns in our WISH assay.

The variation in expression of these genes, despite all being implicated as POM, suggests to us a lack of uniformity within the population. As such, we next sought to examine whether POM cells exhibit differences in behavior during AS colonization.

### POM cells display distinct targeting patterns during early AS colonization

Key to our understanding of AS formation will be the awareness of when and how POM cells colonize. To begin characterizing this process, we first took advantage of available transgenic lines known to label POM: *Tg[Foxc1b:GFP], Tg[Pitx2:GFP], Tg[Lmx1b.1:GFP]*, or NCC: *Tg[FoxD3:GFP], Tg[Sox10:GFP] (12, 31, 33, 34)*. These reporter lines enable single cell distribution analysis while also delineating lineage specification. Due to persistence of GFP protein, these lines do not necessarily represent active expression of their reporter driven promoter but do mark the lineage of POM/NCCs that have, at some point, experienced expression of that POM-associated gene. Transgenic embryos were fixed at key AS developmental stages (24, 26, 38, 30, 48, 56, 72hpf) and immunohistochemistry (IHC) was used to detect GFP. 3D confocal images of the AS were collected for each transgenic line at each timepoint (Fig. 1A). 3D image rendering was employed to subsequently quantify distribution of the cells within the AS (Fig. 1B, Fig. S2).

**Figure 1:**
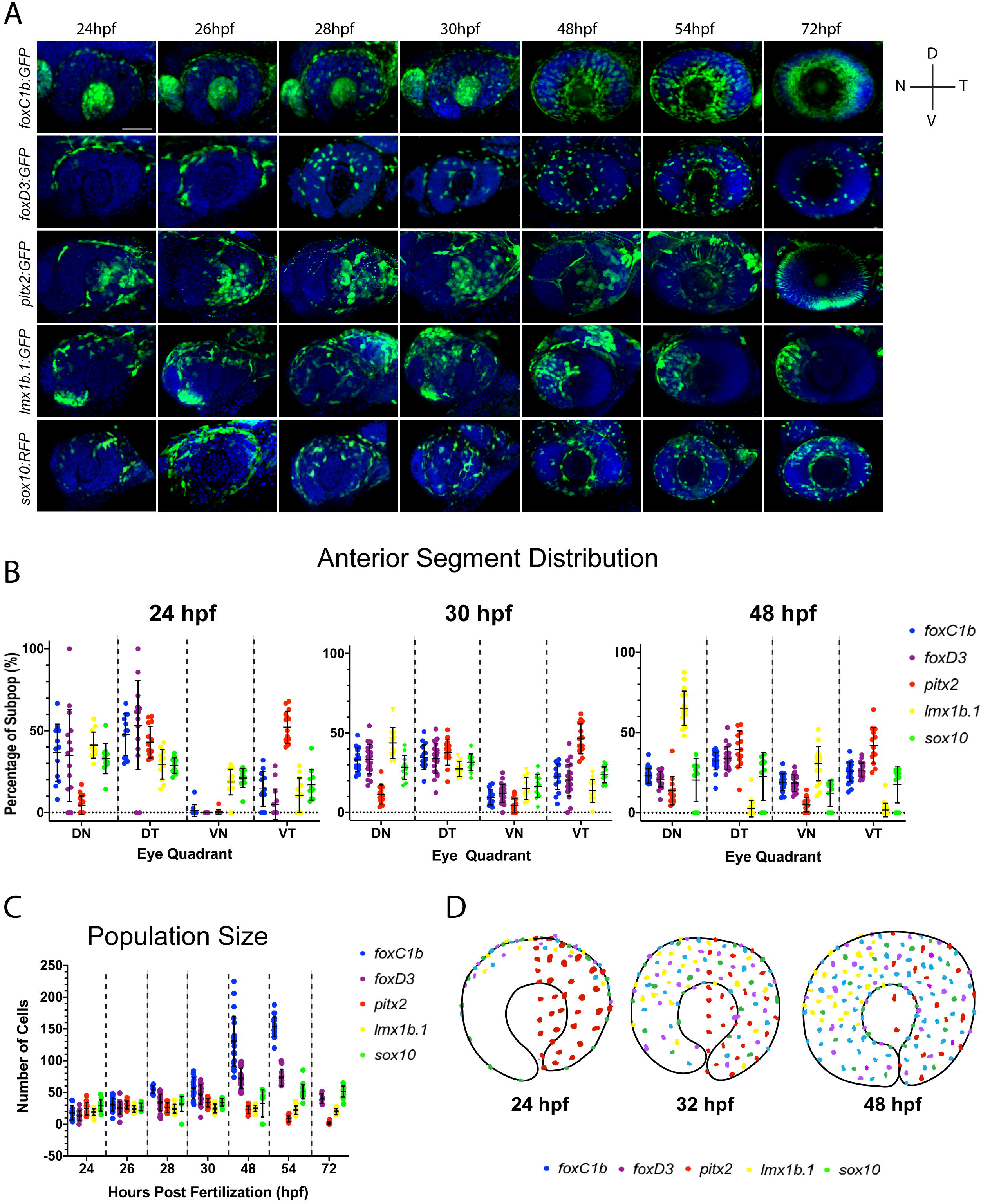
POM subpopulation distribution analysis. **A)** 3D rendering of confocal stacks encompassing the AS in POM transgenic lines between 24-72hpf. GFP+ cells are green, DNA was stained with DAPI (blue). Scale bar=50μm. **B)** Distribution of quantified GFP+ cells within each eye quadrant at 24, 30 and 48hpf. DN=dorsal-nasal, DT=dorsal-temporal, VN=ventral-nasal, VT=ventral-temporal. Distribution is represented as an average percentage of the total number of GFP+ in the AS for each line. **C)** Average cell population size at 24-72hpf. **D)** Model of POM colonization at 24, 30, and 48hpf.

Our assay revealed that initial colonization of the AS begins at approximately 22hpf (data not shown) with GFP+ cells of the *pitx2*-derived population occupying the temporal AS regions (Fig. 1A). At 24hpf *foxc1b, foxD3*, and *lmx1b. 1* derived GFP+ cells have begun to enter the AS across the dorsal most periocular regions, while *sox10*-derived cells begin to occupy all four regions. By 28hpf all of the reporter lines, with the exception of *pitx2*, exhibit GFP+ cells primarily in the dorsal half of the AS. *Foxc1b, foxd3* and *sox10* derived cells continue to spread to the ventral regions with roughly equal distribution throughout the AS by 48hpf. Conversely, *pitx2*-derived cells remain exclusively associated with the temporal half of the AS while *lmx1b. 1*-derived cells gradually re-distribute to occupy the nasal half of the AS. Starting at 54hpf and continuing to 72hpf, Tg[*pitx2*:GFP] expression dissipates from POM cells and initiates in what is likely photoreceptor progenitor cells. Our observations are validated by quantification of each GFP+ population (Fig. 1B). At 24hpf the majority of POM cells are located within the dorsal half of the AS, with the exception of *pitx2*-derived cells. By 30hpf we note a significant reduction in the proportion of *foxc1b, foxD3* and *lmx1b. 1*-derived cells in the dorsal half combined with significant increase of these cells in the ventral half. At 48hpf, we noted equal distribution of *foxc1b, foxd3* and *sox10* derived cells within all regions of the AS, while *lmx1b. 1*-derived cells become predominantly associated with the nasal half of the AS.

A distinct fluctuation in the total number of GFP+ cells within each transgenic line examined throughout the time course was also noted. While *lmx1b. 1, pitx2, and sox10*-derived populations maintained a relatively consistent total cell count, *foxC1b and foxD3*-derived populations grew over time (Fig. 1C). This is particularly evident in the *foxC1b*-population which grew so significantly by 72hpf that it was no longer quantifiable by our assay.

When combining all of our POM AS distribution data, we propose a progressive colonization model (Fig. 1D). In our model, the majority of POM cells colonize the AS in a dorsal-ventral pattern, while pitx2-derived cells uniquely colonize in a temporal-nasal pattern. By 48hpf AS associated POM are equally distributed throughout the four quadrants, except for *lmx1b. 1*- and *pitx2*-derived cells. These POM cells remain uniquely associated with nasal and temporal regions, respectively. Lastly, based on this model we propose that AS-associated POM, which we term anterior segment mesenchyme (ASM) is not a homogenous population and comprises several distinct subpopulations.

### ASM subpopulations exhibit unique migration behavior

The ability to migrate long distances and respond to specific cues is a crucial and well documented behavior of NCC. Cranial NCCs migrate without designated leader or follower identities, instead maintaining a large homogenous population wherein each member exhibits the same migratory capabilities as its neighbors (35–37). We aimed to catalogue the migratory behaviors of ASM cells to determine if they behaved in a similar fashion to cranial NCCs. In particular, we tracked their migration within the AS using *in vivo* 4D imaging. Using this approach, we documented migration of *foxC1b:GFP, foxD3:GFP, pitx2:GFP, lmx1b.1:GFP* and *sox10:RFP*-derived cells (Fig. 2A). Qualitative examination of our data indicated that in all the transgenic lines, ASM cells migrated in a stochastic manner. These cells exhibited seemingly random migration paths, but with a common goal of spreading throughout the AS. This suggests that similar to NCCs, ASM cells lack leader/follower cell identities or chain migration behavior (Fig. 2A, Supplemental Movie 1-5). Careful examination of our data also revealed that *foxC1b, foxd3* and *sox10*-derived populations exhibit frequent cell division while populating the AS. Little to no proliferation was documented in *pitx2* or *lmx1b. 1*-derived ASM subpopulations. Curiously, both *pitx2* and *lmx1b.1*-derived subpopulations also display very specific distributions within the AS (Fig. 2B), while *foxc1b, foxd3* and *sox10* have more homogenous distribution patterns.

**Figure 2:**
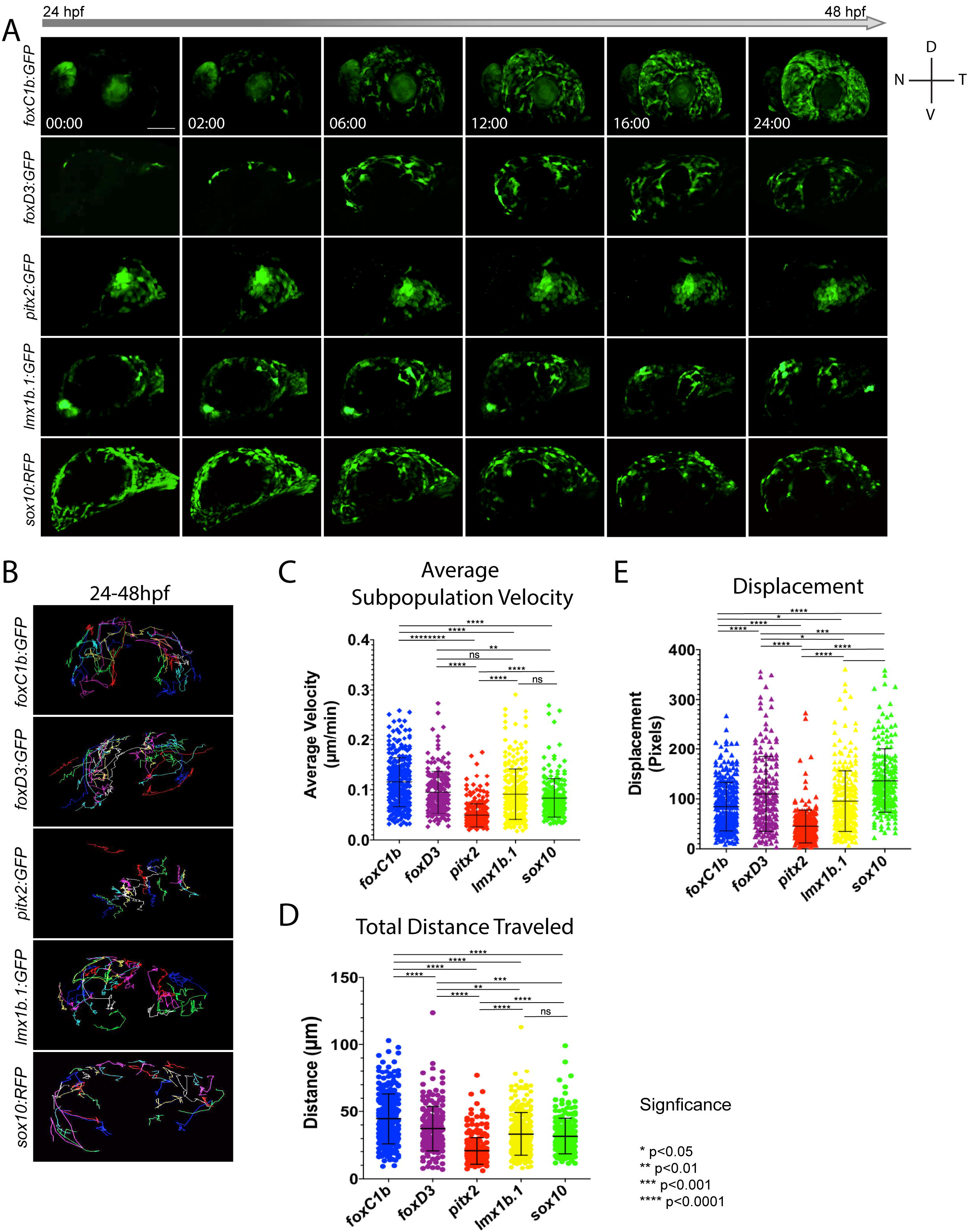
*In vivo* 4D imaging of POM anterior segment colonization. **A)** 4D *in vivo* imaging of the AS conducted between 24-48hpf using POM transgenic lines. Time stamp hours:minutes. Scale bar=50μm. **B)** Individual cell tracking for each transgenic line reveals migratory patterns during early AS colonization. **C-E)** Cell tracking measurements of average migratory velocity (ANOVA p<0.0001), total migration distance (ANOVA p<0.0001), and migratory displacement within the AS. (ANOVA p<0.0001).

To analyze individual migratory behavior, we performed individual cell tracking. As expected, *foxC1b:GFP, foxD3:GFP, lmx1b.1:GFP* and *sox10:GFP*-derived cells migrated in a general dorsal to ventral pattern (Fig. 3B, Supplemental Movie 6-10). *Pitx2:GFP* cells migrated in a generally temporal to nasal direction (Fig. 2B). Tracked cells were analyzed for their total distance traveled, overall displacement, and velocity. *FoxC1b*-derived ASM had the highest velocities (0.115 +/- 0.048μm/min) while the cells of the *Pitx2*-derived subpopulation were the slowest (0.049 +/- 0.023μm/min) (Fig. 2C). *FoxD3, lmx1b. 1*, and *sox10*-derived ASM cells had similar velocities (0.095 +/- 0.042μm/min; 0.091 +/- 0.050μm/min; 0.079 +/- 0.033μm/min). When examining total distance travelled, *foxC1b*-derived cells displayed the farthest distances overall (44.667 +/- 18.531μm), while *pitx2*-derived cells exhibited the shortest (20.719 +/- 9.936μm) (Fig. 2D). *FoxD3, lmx1b.1*, and *sox10*-derived ASM all exhibited similar overall distances traveled (37.339 +/- 16.769μm; 33.267 +/- 15.869μm; 31.688 +/- 13.289μm). Differences in total migratory distance and velocity based on the eye quadrant of origin were also considered. We suspected that ASM cells originating in the dorsal regions of the AS may demonstrate distances and velocities higher than those originating within the ventral regions. However, only cells of the *foxC1b* subpopulation originating in the dorsal temporal region exhibited significant differences in migration parameters from the overall population. These cells travelled shorter total distances and had slower velocities when compared to the other three quadrants (Fig. S3).

**Figure 3:**
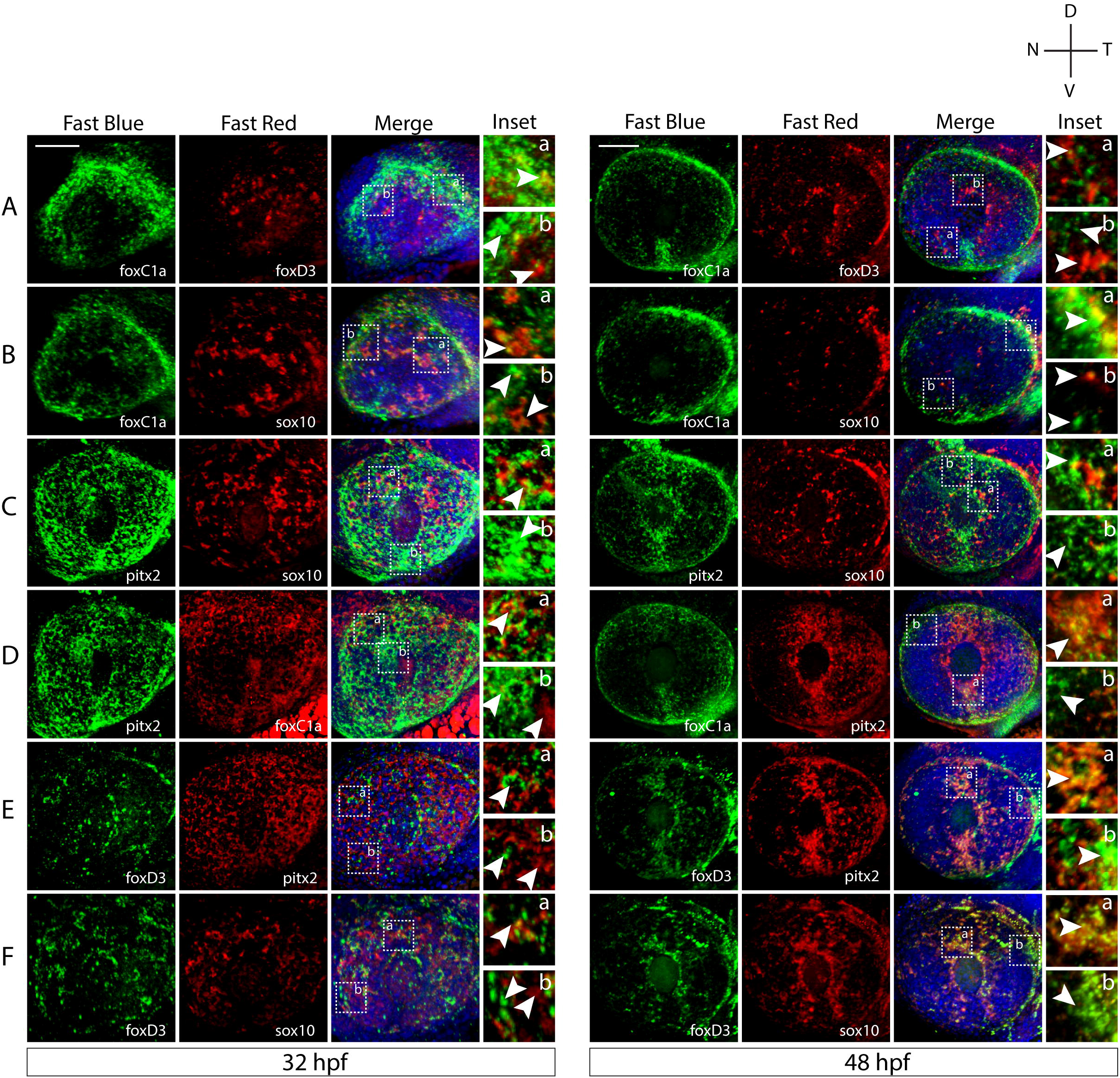
Two-color fluorescent *in situ* hybridization supports ASM heterogeneity. **A-F)** Two-color-fluorescent WISH (FWISH) performed for all possible combinations of *foxC1a, foxD3, pitx2* and *sox10* at 32 and 48hpf. DAPI is in blue. White arrows within inset panels (dashed squares) display instances of individual (b) and co-expression (a). Scale bar=50μm.

Lastly, we measured the degree of directed migration by examining displacement within the AS (Fig. 2E). Cells within the *sox10* subpopulation showed the highest overall displacement (136.885 +/- 63.327pixels), followed by cells in the *foxD3* subpopulation (109.70 +/- 75.434pixels). Displacement of *foxC1b* and *lmx1b. 1* was found to be 84.632 +/- 49.223pixels and 95.559 +/- 61.285pixels, respectively. Similar to our previous observations for velocity and distance, the *pitx2* subpopulation exhibited the least amount of displacement (44.698 +/- 33.008pixels). These data suggest, that cells in the *sox10*- and *foxD3*-derived subpopulations, associated with NCC identity (25, 27, 28), respond or achieve more effectively to directed or targeted migration. Cells associated with a more traditional POM identity, *foxC1b-, lmx1b. 1*- and *pitx2*-derived (15, 18), appear to have more stochastic migration paths.

Taken together, while all of the ASM subpopulations examined migrate in a largely stochastic, NCC-like manner, their migratory mechanics and targeting behavior significantly differ. These conclusions further support our hypothesis that the ASM is a heterogenous population.

### Co-expression analysis confirms anterior segment mesenchyme heterogeneity

Having observed unique distributions and migratory behavior within the ASM, we next examined if there was co-expression of POM marker genes amongst the subpopulations. To study these relationships, we performed two-color-fluorescent *whole mount in situ* hybridization (FWISH) at 32 and 48hpf. We focused our attention to the expression of *foxc1a, foxd3, sox10* and *pitx2* as they represent our transgenic reporter lines (Fig. 1). 3D confocal imaging qualitatively indicated that all these POM markers clearly exhibit both overlapping and individualized expression patterns (Fig. 3). *Pitx2* exhibited broad expression throughout the AS and a high degree of co-expression with all the other markers (Fig. 3C-E). *FoxC1a* was expressed in the outer lateral regions of the AS, but mostly absent from the center. Also, *foxC1a* exhibited a high degree of co-expression with *pitx2* and partial co-expression with *foxD3* and *sox10* (Fig. 3A,B,D). The expression of *sox10* was most pronounced in the dorsal AS and had the highest degree of co-expression with foxD3 (Fig. 3B,C,F). *FoxD3* expression was detected throughout the AS and exhibited a slight degree of co-expression with the other markers (Fig 3A,E,F). We also analyzed expression of *eya2* and observed a high degree of co-expression with *pitx2* and slight co-expression with all the other markers (Fig. S4).

According to our results, ASM co-expression occurred among almost all different combinations of POM markers but varied in degree and localization within the AS. *Sox10* and *foxD3*, both NCC markers, appear to be co-expressed in multiple cells distributed over the entire AS at 32hpf. Co-expression of *pitx2* and *foxc1* is limited to the lateral area of the AS while co-expression of *pitx2* and *eya2* occurs throughout the AS. Although we do see a degree of co-localization amongst all of POM marker genes, the distinction between each gene’s expression pattern remains evident. This suggests that individual ASM cells exhibit a range of POM marker expression patterns which also supports our hypothesis that ASM is heterogenous during AS development.

### ASM subpopulations exhibit differing transcriptomic expression patterns

Having confirmed the subdivision of the AS associated POM into several ASM subpopulations, our final goal was to complete a transcriptomic comparison of the groups. To do so, we isolated GFP+ ASM populations from cranium of 48hpf transgenic embryos via FACS and subsequently sequenced total RNA. We generated transcriptomes from *foxc1b, foxd3, pitx2*, and *sox10*-derived ASM and performed a 4-way expression analysis using RSEM and RStudio (Table 2).

When compared to the other three populations, the *sox10* expressing subpopulation had the most unique signature, containing 700 genes which were differentially expressed (Fig. 4A). This is likely based on their ability to maintain NCC’s multipotent nature. *FoxD3* expressing cells, contained only 40 transcripts unique to this subpopulation, possibly owing to the fact that *foxD3* is also a marker of NCCs and likely shares expression profiles with cells expressing *sox10* (Fig. 4C). *FoxC1b* expressing cells had the most unique expression pattern of the ASM populations with 400 differentially expressed transcripts, suggesting that it is the most unique subpopulation within the ASM sub-groups examined to date (Fig. 4B). This may be indicative of the *foxC1b* subpopulation having broad regulatory control over the other, smaller subpopulations. The most seemingly redundant subpopulation is that of the *pitx2* expressing cells with only 12 transcripts having a unique expression pattern (Fig. 4D). *Pitx2* has been heavily implicated in ASD, particularly Axenfeld-Rieger syndrome, making its relative lack of individuality surprising (10, 11, 14, 17). This data further supports our hypothesis that the AS is colonized by several ASM subpopulations. It stands to reason that these subpopulations may eventually give rise to specific AS structures.

**Figure 4:**
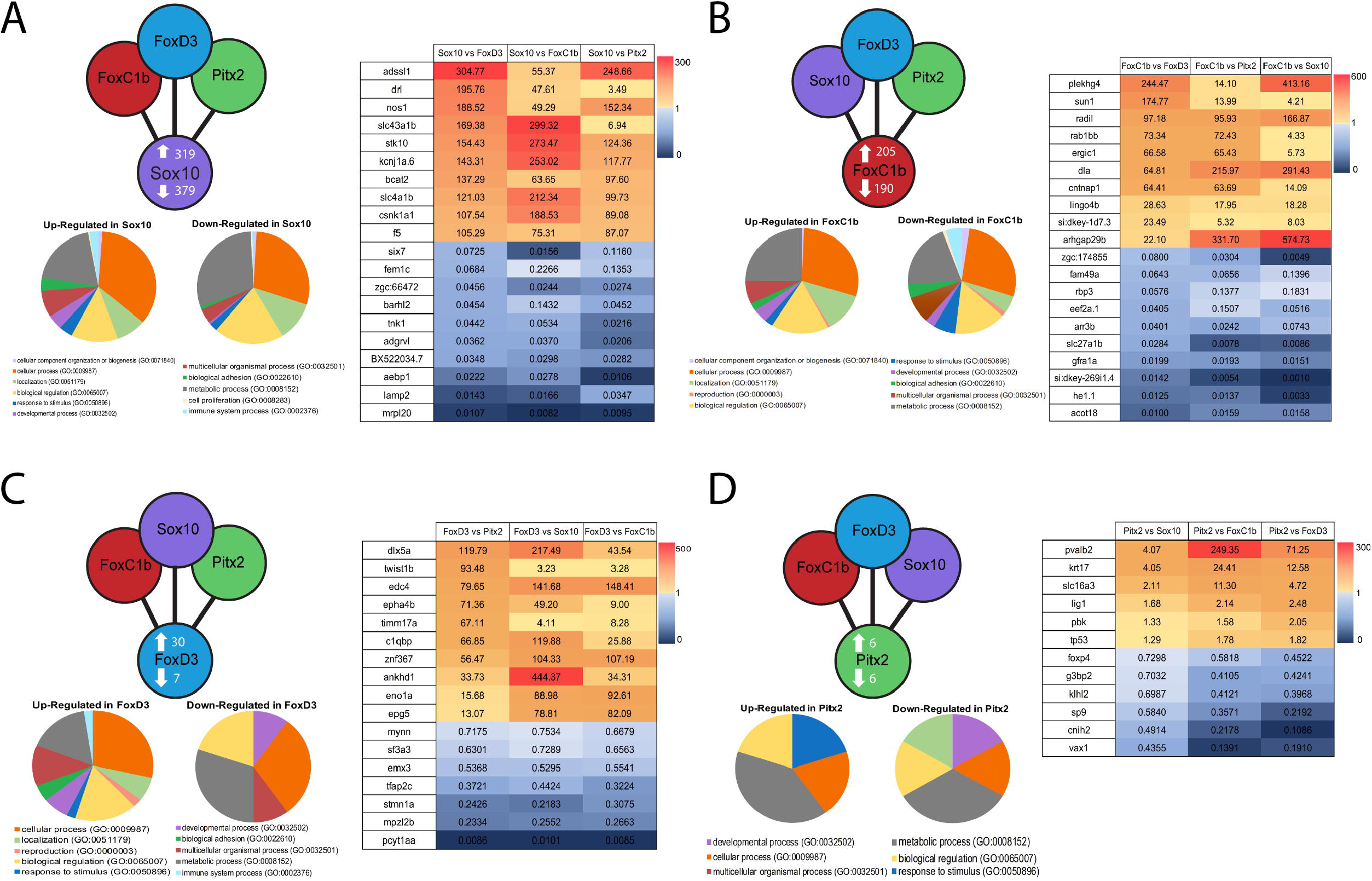
Four-way transcriptomic analysis of POM populations at 48hpf. Transcriptomic profiles of each subpopulation compared to the other three. Counterclockwise: the total number of up and down-regulated genes, gene ontology of all differentially expressed genes, and a heat map of top 10 up and down-regulated genes. **A)** *sox10* profile **B)** *foxC1b* profile. **C)** *foxD3* profile **D)** *pitx2* profile.

In conclusion, our data indicate that POM cells targeting to the anterior segment, for which we suggest the term ASM, are not homogenous. Rather, the ASM comprises of several subpopulations. We have characterized four ASM subpopulations each with distinct distributions, population sizes, migratory velocities, displacements and transcriptional profiles. Our findings open a wide range of possible new investigative paths in the area of AS structure development and ASD disorders. Including a more complete understanding of not only single gene involvement, but rather subpopulation influence on one another.

## MATERIALS AND METHODS

### Zebrafish maintenance

Zebrafish lines were bred and maintained in accordance with IACUC regulations at the University of Kentucky (protocol # 2015-1370). AB strain was used as wildtype. Transgenic lines used were: Tg[*foxC1b:GFP*] (Dr. Bryan Link), Tg[*foxD3:GFP*] (Dr. Lister), Tg[*pitx2C4:GFP*] (Dr. Elena Semina), and *Tg[lmx1b.1:GFP]* (Dr. Bryan Link), *Tg[sox10:RFP]* (Dr. Lister). All embryos were raised for the first 24 hours post fertilization in embryo media (E3) at 28°C. After 24 hours, E3 media was replaced with embryo media containing 1-phenyl 2-thiourea (PTU) every 24 hours to maintain embryo transparency.

### Whole-mount in situ hybridization (WISH)

Whole-mount in situ hybridization was performed on a minimum of 50 embryos for each time point (12, 18, 24, 32, 48 and 72hpf) and for each probe used. DIG and FITC labeled RNA probes were generated using PCR incorporating T7 promoters in the primers and ultimately transcribed with T7 polymerase (Roche). Forward and Reverse primer sequences are listed in Supplemental Table 1. WISH protocol was performed as previously described (38). Dorsal, lateral, and ventral images of embryos were captured using a Nikon Digital Sight DS-U3 camera and Elements software. Images were adjusted for brightness using Adobe Photoshop.

### Immunohistochemistry (IHC) and Distribution Analysis

Approximately 30 embryos were imaged for each transgenic line at each of the given time points (24, 26, 28, 30, 48, 54, and 72hpf). Embryos were fixed overnight at 4°C using 4% PFA. PFA was washed out with PBST 4 times for 5 minutes each. Embryos were permeabilized with Proteinase K (10μg/ml) at the following times (24hpf = 5mins; 26hpf = 6mins; 28hpf = 7 mins; 30hpf = 9 mins; 48hpf = 20 mins; 54hpf = 25mins; 72hpf = 40mins), washed with PBST and then blocked with 5% goat serum (1g/100ml), 1% BSA in a solution of 1x PBST for at least 2 hours at room temperature. Primary antibody (Rockland rabbit anti-GFP) was diluted at 1/200 in blocking buffer and incubated overnight at 4°C on rotation. The following day, the primary antibody solution was washed out with PBST 5 times for 15 minutes each. Secondary antibody (Alexa Fluor 488 anti rabbit, 1/1000) and DAPI (1/2500) were diluted in blocking buffer and incubated for 1 hour on rotation in the dark at room temperature. Embryos were washed 2x for 15 minutes with PBST in the dark.

After staining, embryos were embedded in a 1.2% Low-gelling agarose in a 1-inch glass bottom cell culture dish (Fluordish, World Precision Instruments) and visualized using a Nikon C2+ confocal microscope with a 20x (0.95NA) oil immersion objective. The anterior segment of the eye was imaged in 3D in the lateral position as a 100μm z-stack using 3.50μm steps. All images were captured using Nikon Elements software and adjusted using Adobe Photoshop. Images generated from IHC analysis were rendered in 3D using Nikon Elements Viewer software. Eyes were divided into 4 quadrants: dorsal nasal, dorsal temporal, ventral nasal, and ventral temporal (Figure S2). Nasal and temporal regions were divided by a vertical straight line through the center of the lens, while dorsal and ventral were divided by a horizontal straight line through the center of the lens. For distribution analysis, GFP positive cells were manually counted based on their position within one of the 4 quadrants of a 3D constructed anterior segments. For each timepoint 25+ embryos from 2-3 independent trials were imaged for quantification.

### Two Color Fluorescent WISH

RNA probes were generated using the MEGAscript T7 transcription Kit (Ambion) in combination with 10x RNA labeling mixes for both DIG and FITC. Double *in situ* hybridization was performed according to the protocol by (39). This included exposing embryos to acidified methanol and adding Dextran Sulfate into the hybridization reaction. Staining was done by combining Fast Blue and Fast Red dyes (10μL/1mL). After successful *in situ* double staining, embryos were additionally stained using DAPI (1μL/1mL) and imaged using a NIKON C2+ confocal microscope. Images were adjusted using Adobe Photoshop. 15-20 embryos were analyzed for each probe combination.

### Time-lapse confocal in vivo Imaging

Embryos from each of the previously mentioned transgenic lines were collected and raised in E3 media at 28°C. Fluorescent embryos were placed in E3 PTU media including 3-amino benzoic acidethylester (Tricaine) to prevent pigmentation and anesthetize them, respectfully. They were then dechorinated and embedded laterally in 1% low gelling agarose in a 1-inch glass bottom cell culture dish (Fluordish, World Precision instruments). Real-time imaging was conducted at 28°C using a Nikon C2+ confocal microscope and a 20x (0.95NA) oil immersion objective. 3D z-stacks over a 75μm thickness with a slice size of 3.5μm were collected to encompass the entire developing anterior segment. Z-stack images were taken at 10 minute intervals over a 24 hours period (embryos imaged: n=12 *foxC1b:GFP*, n=9 *foxD3:GFP*, n=13 *pitx2:GFP*, n=10 *lmx1b. 1:GFP*, n=7 *sox10:RFP*). Data were collected and rendered using Nikon Elements software.

### Cell Migration Tracking and Displacement Analysis

Completed 4D live imaging files were uploaded into FIJI software for analysis. Approximately 25 cells were manually tracked per video file. Tracked cells were measured for total distance traveled (μm), average velocity (μm/min), and total displacement. Tracked cells were randomly selected from all four eye quadrants to ensure all eye regions were represented, as well as all time frames. After tracking, data was exported to Microsoft Excel for statistical analysis. Cell displacement was measured manually using FIJI software and statistically analyzed in Microsoft Excel and Graphpad Prism8.

### Fluorescence Activated Cell Sorting (FACS) and RNA extraction

Embryos from each of the POM subpopulation transgenic lines were dechorianted and incubated in E3 media at 28°C until 48hpf. At this time, embryos were anesthetized using 3-amino benzoic acidethylester (Tricaine) and decapitated just posterior of the eyes. Heads were collected and incubated for 2 minutes in 0.25% Trypsin + EDTA at 37°C. After incubation, a 20G needle and syringe were used to dissociate the tissue before the tube was placed back at 37°C for 2min. This process was repeated 4 times. After incubation, the dissociated cells were strained using a 40μm filter (VWR) and spun down for 10mins at 3500rpm at 4°C. The supernatant was removed and the pellet resuspended in 1x PBS + 2 mM EDTA. Cells were sorted for GFP+ cells at the University of Kentucky Flow Cytometry and Immune Monitoring Core at the Markey Cancer Center. ∼250,000-500,000 cells were collected in three distinct trials. Following cell sorting, RNA was extracted using Trizol according to the manufacturers protocol. Concentration and purity of extracted RNA was performed using the Bioanalyzer at the University of Kentucky HealthCare Genomics Core at the UK Albert B. Chandler Hospital. RNA was then DNAse treated (TURBO DNAfree Kit, Thermo Fisher) and sent to Applied Biological Materials (ABM) for Illumina sequencing. 20-40 million 150bp paired-end reads were generated from each of the 3 samples/transgenic line examined.

### Bioinformatics

Once sequencing results were obtained, bioinformatic analysis was completed using RSEM. Transcripts which were differentially expressed between each of the four subpopulations independently with a confidence interval of 95% or greater were considered significant. Subpopulation sequencing was then compared to find patterns showing gene expression profiles unique to each of the four subpopulations using RStudio. Genes showing unique signatures in a specific subpopulation were compiled with the RSEM data to generate statistically significant expression patterns for each of the subpopulations. Gene ontology charts and heat maps were generated using Microsoft Excel.

### Statistics

One-way ANOVA analysis and Unpaired *t*-tests were performed using Microsoft Excel and GraphPad Prism8 software. All graphs are shown with their respective means and standard deviations. Values were considered significant by the conventional standard: *P* value of 0.05 or less.

## SUPPLEMENTARY FIGURES

**Supp Figure 1: WISH for previously described POM and neural crest-related genes**.

POM mRNA expression patterns during early to late stage eye development in the lateral view. POM genes included *foxC1a, foxC1b, foxD3, pitx2, lmx1b.1, lmx1b.2*, and *sox10*. Expression was analyzed at key time points from 12-72hpf (additional POM-related in situ characterizations can be found in the supplementary materials). The expression of most POM genes is visible, although not guaranteed to be within the dorsal regions. All genes tested displayed expression around the optic cup by 24hpf. *Foxc1, pitx2*, and *sox10* in particular show strong expression surrounding the optic cup and on the surface of the anterior segment from 24-72hpf. *Lmx1b. 1* and *lmx1b.2* do not show strong expression directly surrounding the eye despite its implications in anterior segment development, suggesting a cell nonautonomous mechanism. Expression later appears within the anterior segment at 48-72hpf.

**Supp Figure 2: AS eye quadrant quantification scheme**.

The eye was split evenly through the lens horizontally and vertically to generate four distinct quadrants.

**Supp Figure 3: *FoxC1b* cell distances traveled and velocity of migration varies by eye quadrant of origin**.

In order to determine if the quadrant of origin had an effect on the distance or velocity of migrating cells within a subpopulation, FIJI migration data was analyzed based on the starting position of a cell on the AS. Cells originating in the dorsal temporal region of the *FoxC1b* subpopulation migrated a shorter distance (**A**) and at a slower velocity (**B**) than cells originating in the other three quadrants.

**Supp Figure 4: Anterior segment ASM heterogeneity for *eya2* expression**.

**A-F)** Two-color-fluorescent WISH (FWISH) performed for all possible combinations of *eya2 VS foxc1a, foxD3, pitx2* and *sox10* at 32 and 48hpf. DAPI is in blue. White arrows within inset panels (dashed squares) display instances of individual (b) and co-expression (a). Scale bar=50μm.

**Supp Table 1: *WISH* Primer Sequences**. mRNA forward and reverse primer sequences for all POM and NCC-related genes.

**Supp Table 2: 4-way transcriptomic analysis**. Results of fold change between the four ASM populations analyzed.

**Movie 1: *FoxC1b:GFP* 4D imaging (24-28hpf)**.

**Movie 2: *FoxD3:GFP* 4D imaging (24-48hpf)**.

**Movie 3: *Pitx2:GFP* 4D imaging (22-46hpf)**.

**Movie 4: *Lmx1b.1:GFP* 4D imaging (24-48hpf)**.

**Movie 5: *Sox10:RFP* 4D imaging (23-47hpf)**.

**Movie 6: *FoxC1b:GFP* Tracking analysis**

**Movie 7: *FoxD3:GFP* Tracking analysis**.

**Movie 8: *Pitx2:GFP* Tracking analysis**.

**Movie 9: *Lmx1b.1* Tracking analysis**.

**Movie 10: *Sox10:RFP* Tracking analysis**.

